# Isolation and Identification of Enterovirus D111 in a Child with Acute Flaccid Paralysis in Nigeria

**DOI:** 10.1101/479873

**Authors:** T.O.C. Faleye, M.O. Adewumi, O.T. Olayinka, J.A Adeniji

**Author notes:** Corresponding Author: Faleye T.O.C.; 08023840394.

## Abstract

The WHO recommended cell-culture-based algorithm requires enterovirus (EV) isolates to produce reproducible cytopathic effect (R-CPE) in RD and/or L20b cell lines. Samples with non-reproducible CPE (NR-CPE) are considered negative for EVs. We investigated whether there could be EVs lurking in samples with NR-CPE.

Fifty-nine (59) cell culture supernatants (CCS) (collected between 2016 and 2017) recovered from RD and L20b cell culture tubes with NR-CPE, were analyzed in this study. The tubes had been previously inoculated with stool suspension from children (<15 years) in Nigeria with acute flaccid paralysis (AFP). All CCS were screened for Enteric Adenoviruses and group A Rotavirus using a rapid immunochromatographic test kit. Subsequently, they were passaged in HEp-2 cell line. All isolates were subjected to RNA extraction, cDNA synthesis, three (5^*l*^-UTR, VP1 and EV Species C [EV-C]) different PCR assays and sequencing of amplicons. EVs were further subjected to Illumina sequencing.

All CCS were negative for Adenoviruses and group A Rotaviruses. Four CCS produced R-CPE in HEp-2 cell line, three of which were positive for the 5^*l*^-UTR assay. Of the 3 isolates two and none were positive for the VP1 and EV-C assays, respectively. One of the two VP1 amplicons was successfully sequenced and identified as Echovirus 1(E1). Illumina sequencing of the three 5^*l*^- UTR positive isolates confirmed the E1 isolate and typed the remaining two as EV-Ds (94 and 111).

We describe the first EV-D94 and 111 isolates of Nigerian origin. We also show that NR-CPE could sometimes be caused by EVs that do not produce R-CPE in RD and L20b cell lines but do so in other cell lines like HEp-2.

## INTRODUTION

There are 15 Species in the genus Enterovirus, family *Picornaviridae*, order *Picornavirales* (http://www.picornaviridae.com/enterovirus/enterovirus.htm). Enterovirus Species C (EV-C) remains the type Species and Poliovirus (PV), the prototype member of the genus. Enteroviruses (EVs) have a naked capsid with icosahedral symmetry and a diameter of 28-30nM. The capsid encapsulates a positive sense, single-stranded RNA genome that is 7,500 nucleotides in length. The genome has one open reading frame (ORF) that is flanked by untranslated regions on both ends. The ORF encodes one polyprotein that is autocatalytically cleaved to make 11 proteins; four structural proteins (VP1 – VP4) and seven non-structural proteins (2A-2C and 3A-3D). Classically, EV identification was done by neutralization assays. However, since a correlation was shown to exist between serological types and VP1 sequence (Oberste et al., 1999), VP1 amplification and sequencing became the standard for EV identification. Recently, the European Non-polio Enterovirus Network (ENPEN) recommended (Harvala et al., 2018) that in cases where VP1 data (or the full genome) is not available, VP2 and VP4 can be utilized for EV identification.

Most of the EV data available globally is courtesy the Global Polio Laboratory Network (GPLN); the laboratory arm of the Global Polio Eradication Initiative (GPEI). GPEI was established to eradicate PV globally and have so far reduced the incidence of poliomyelitis by >99.9% (Kew and Pallansch, 2018). Their approach has so far included a blend of surveillance for PV (in acute flaccid paralysis [AFP] cases as well as in the environment) and vaccination (using both inactivated and live attenuated poliovirus vaccines) to interrupt PV transmission chains (Kew and Pallansch, 2018).

In the GPLN, PV detection is done using a cell culture-based algorithm that requires PV isolates to produce reproducible cytopathic effect (R-CPE) in RD and L20b cell lines (WHO, 2003, 2004, 2015). The algorithm also results in the isolation of non-polio enteroviruses (NPEVs) which are also required to produce R-CPE in RD and/or L20b cell lines (WHO, 2003, 2004, 2015). Samples with non-reproducible CPE (NR-CPE) are considered negative for EVs. This phenomenon of NR- CPE has been suggested to be caused by nonspecific cytotoxicity, Adenovirus or Reoviruses (to which the Rotaviruses belong) (WHO, 2007). However, we (Adeniji et al., 2018) recently showed that EVs could also contribute to NR-CPE. In this study, we go a step further to find out how often the three (Adenoviruses, group A Rotaviruses and EVs) virus types are present in cell culture supernatants (CCS) recovered from L20b cell culture tubes with NR-CPE.

## METHODS

### Samples

Fifty-nine (59) cell culture supernatants (CCS) were analyzed in this study following the algorithm depicted in Figure 1. All 59 CCS were recovered from L20b cell culture tubes with NR-CPE. The tubes had been previously inoculated with stool suspension from children (≤15 years old) in Nigeria with acute flaccid paralysis (AFP). Prior this study, the samples were accumulated over the course of one (1) year (2016-2017) during which they were stored at −20°C

**Figure 1:**
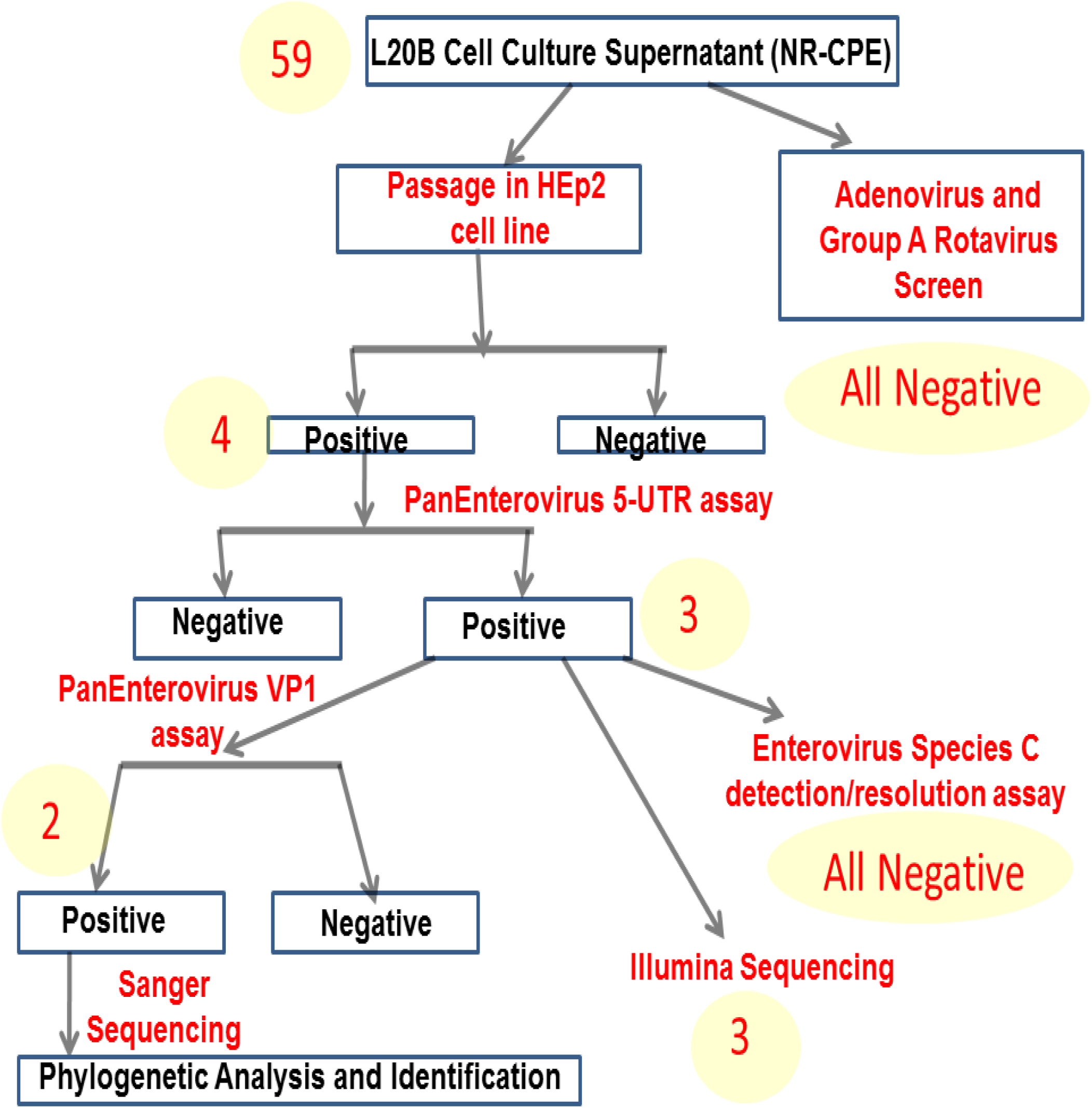
A schematic representation of the algorithm followed in this study.

### Adenovirus and Group A Rotavirus Screen

A rapid immunochromatographic test kit (Rotavirus Group A antigen/Enteric Adenovirus Antigen Rapid test kit, Accumed Technology Co. Ltd, Beijing) was used to screen the samples for Enteric Adenoviruses and Group A Rotavirus. As controls, a fecal suspension from a child with Rotavirus diarrhea that we had confirmed by molecular testing (unpublished data) and reference Adenovirus 1 and 6 from the BBSRC (a kind gift from Oladipo K.E.) were used as positive controls.

### Passage in Hep2 Cell line

The HEp-2 cell line used in this study was maintained in Dulbecco’s Modified Eagle Medium (DMEM) supplemented with 2% Fetal Calf Serum, 100IU/ml penicillin and 100mg/ml streptomycin. All CCS were passaged in HEp-2 cell line and incubated for 14 days at 37^0^C. For each CCS, 200µL was inoculated into two tissue culture tubes each containing HEp-2 cells. The tubes were then incubated at 37^0^C and monitored for 14 days. Daily the tubes where visually screened (using an inverted microscope) for EV characteristic cytopathic effect (CPE). Tubes in which CPE was observed were further passaged to confirm such. Samples were only declared negative for CPE if none was observed after 14 days of observation.

### Total RNA extraction and cDNA synthesis

Total RNA was extracted from all samples that produced CPE in HEp-2 cell line using the Total RNA extraction kit (Jena Bioscience, Jena, Germany). RNA was extracted following the guidelines detailed by the manufacturer. The SCRIPT cDNA synthesis kit (Jena Bioscience, Jena, Germany) was used for cDNA synthesis following manufacturer’s instructions. Two different cDNAs were prepared; cDNA 1 (using random hexamers, Faleye et al., 2017) and cDNA 2 (using primers AN32 to AN35, Nix et al., 2006).

### Polymerase Chain Reaction (PCR) Assays

Three (PE-5-UTR, EC 3Dpol/3^I^-UTR PCR and PE-VP1) different PCR assays that target three different (5^I^-UTR, 3Dpol-3^I^-UTR and VP1) EV genomic regions were done in this study (Table 1). It is however crucial to mention that the VP1 assay is a semi-nested PCR assay with PE-VP1a and PE-VP1b being the first and second round assays, respectively.

**Table 1:** Details of PCR assays

| S/N | Assay Name | Primer Name | Region Amplified | Amplicon Size (bp) | Extension Time (Seconds) | cDNA used | Reference |
| --- | --- | --- | --- | --- | --- | --- | --- |
| 1 | PE-5-UTR | PanEnt-5'-UTR-F<br>PanEnt-5'-UTR-R | 5'-UTR | ~114 | 30 | 1 | Oberste *et al.*, 2006 |
| 2 | EC 3Dpol/3'-UTR PCR | HEV-C-3D<br>HEV-C-3'-UTR | 3Dpol-3'-UTR | ~429 | 30 | 1 | Adeniji and Faleye, 2014 |
| 3 | *PE-VP1a | 224<br>222 | VP3-VP1 | ~750 | 60 | 2 | Nix et al., 2006 |
| 4 | *PE-VP1b | AN89<br>AN88 | VP1 | ~350 | 30 | NA | Nix et al., 2006 |

NA = Not Applicable
\*Both PE-VP1a and PE-VP1b are both components of the Nix et al., 2006 semi-nested PCR protocol. PE-VP1a is the first round while PE-VP1b is the second round PCR.
cDNA 1= synthesized using random hexamers (Faleye et al., 2017)
cDNA 2= synthesized using Oligos AN32-AN35 (Nix et al., 2006)

All primers used were made in 100µM concentrations. All PCR assays were done in 30µL volumes containing 6µL of Red Load PCR mix (Jena Bioscience, Jena, Germany), 0.3 µL of each of the forward and reverse primers, 5µL of cDNA or first round PCR product and 18.4µL of PCR grade water. Amplification was done using the Veriti thermal cycler as follows; 94°C for 3 min followed by 45 cycles of 94°C for 30s, 42°C for 30s and 60°C for 30s with ramp from 42°C to 60°C adjusted to 40%. This was followed by 72°C for 7 min and held at 4°C till terminated. It should however be noted that the extension temperature varied based on the expected amplicon size as detailed in Table 1. All PCR products were resolved on 2% agarose gels stained with ethidium bromide and viewed using a transilluminator.

### Sanger Sequencing

Only the amplicons generated by the PE-VP1b assay (Table 1) were subjected to Sanger sequencing. The amplicons were shipped to Macrogen Inc, Seoul, South-Korea, where gel extraction and Sanger sequencing were done. Sanger sequencing was done using the primers AN89 and AN88.

### Illumina Sequencing

For Illumina Sequencing, cDNA 1 was used. The genomes were amplified in overlapping fragments of 2 – 3kb as previously described (Faleye et al., 2018a). Irrespective of the intensity of amplicon or whether or not there were amplicons, for each isolate, the overlapping genomic fragments were pooled and shipped to a commercial facility (MR DNA, Texas, USA) where library preparation (Nextera DNA sample preparation kit), Illumina sequencing (paired end, 300 cycles, HiSeq) were done. The Illumina sequencing data was assembled using the Kiki assembler v0.0.9. Subsequently, detection of EV contigs was done using the Enterovirus Genotyping Tool (EGT) v1.0 (Kroneman et al., 2011).

### Enterovirus identification and Nucleotide accession numbers

Enterovirus types were determined using the EGT v1.0 (Kroneman et al., 2011). Nucleotide sequences generated in this study have been submitted to GenBank under the accession numbers MH607128, MK190417-MK190420.

### Phylogenetic Analysis

All the sequences of the EV types of interest in GenBank were downloaded. Using the EGT v1.0 (Kroneman et al., 2011), sequences that did not cover the VP1 region of interest were removed. The sequences were then aligned using the ClustalW program in MEGA 5 software with default settings (Tamura et al., 2011). Subsequently, neighbor-joining trees were constructed using 1000 bootstrap replicates (Tamura et al., 2011). The accession numbers of sequences retrieved from GenBank for analysis are shown in the phylograms.

## RESULTS

### Adenovirus and Group A Rotavirus Screen

All the samples screened in this study were negative for the Enteric Adenovirus and Group A Rotavirus Screen (Figure 1). However, the positive controls (a fecal suspension from a child with Rotavirus diarrhea and reference Adenovirus 1 and 6) were repeatedly and consistently detected by the rapid detection kit.

### Passage in HEp2 Cell line

On passage in HEp2 cell line, only 4 (6.8%) of the 59 samples yielded isolates that were reproducible on passage in HEp2 cell line (Figure 1).

### PCR assays

Three (75%) of the four isolates were positive for the PanEnterovirus 5-UTR (PE-5-UTR) assay. None of the three (3) PE-5-UTR positive isolates was positive for the Enterovirus Species C detection/resolution (EC 3Dpol/3^I^-UTR PCR) assay. Two (2) of the three (66.7%) PE-5-UTR positive isolates were positive for the PanEnterovirus VP1 (PE-VP1) assay (Figure 1).

### Sanger Sequencing

Of the two (2) samples positive for the PE-VP1 and shipped to the sequencing facility, only one was successfully sequenced. The other could not be sequenced due to insufficient volume. The sequenced isolate was typed as Echovirus 1 using the EGT (Table 2).

**Table 2:** Identity of PE-5UTR positive samples using VP1 Sanger and Illumina data

| EV Isolate S/N | PE-VP1 Assay | IDENTITY |  | EV Species |
| --- | --- | --- | --- | --- |
|  |  | Sanger | Illumina |  |
| 1 | Positive | Not Sequenced | EV-D94 | EV-D |
| 2 | Positive | Echovirus 1 | Echovirus 1 | EV-B |
| 3 | Negative | Not Applicable | EV-D111 | EV-D |

### Illumina Sequencing

This showed that isolates 1, 2 and 3 contained EV-D94, E1 and EV-D111 (Table 2). Two contigs were assembled for each of isolates 1 and 3 while only one was assembled for isolate 2 (Figure 2).

**Figure 2:**
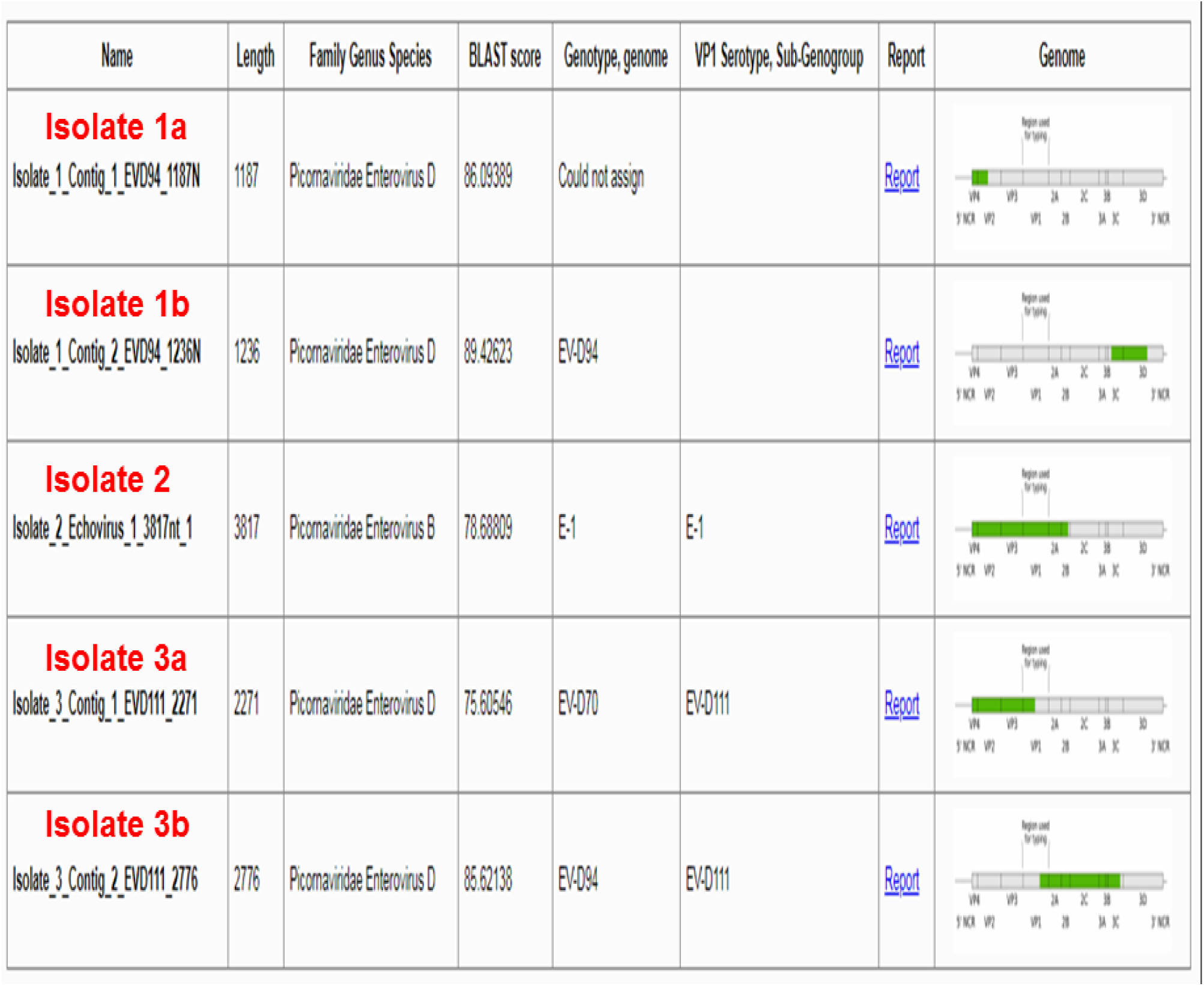
The Contigs assembled from the Illumina sequencing data generated for the three PE- 5UTR positive isolates. The contigs and the genomic regions covered are displayed using the EGT.

### A Glitch in the Enterovirus Genotyping Tool

It was observed that for isolate 3 (Table 2), there was disparity in the genotype and serotype as determined by the EGT (Figure 2). In fact, there was also variation from one contig of isolate 3 to the other. To resolve this aberration and confirm the identity of isolate 3, nucleotide similarity between the VP1 sequence of isolate 3 (and other EV-D111 from GenBank) and a reference EV- D70 was estimated (Table 3). This confirmed that isolate 3 was indeed EV-D111. Furthermore, using the EGT to type the six (6) EV-D111 VP1 sequences available in GenBank, prior this study, showed that the aberration was not unique to isolate 3 (Figure 3)

**Table 3:** Estimation of similarity between EV-D111 and EV-D70 VP1 to confirm identity of Isolate 3

| **EV-D111** | **Reference EV-D70** | **Distance (%)** | **Similarity (%)** |
| --- | --- | --- | --- |
| JX431302 EVD111 111 TOK-230 HC 2008 | D00820 EVD70 J670 71 | 31.03 | 68.97 |
| JN255684 EVD111 CAF-NGB-06-107 2006 | D00820 EVD70 J670 71 | 32.15 | 67.85 |
| EF127249 EVD111 17-04 DRC 2000 | D00820 EVD70 J670 71 | 33.27 | 66.73 |
| EVD111 NGR 2017 | D00820 EVD70 J670 71 | 33.89 | 66.11 |
| JF416935 EVD111 KK2640 Chimpanzee NHP 2011 | D00820 EVD70 J670 71 | 33.94 | 66.06 |
| JN255683 EVD111 CAF-BAN-06-150 2006 | D00820 EVD70 J670 71 | 34.52 | 65.48 |
| JN255685 EVD111 CAF-OUP-05-059 2005 | D00820 EVD70 J670 71 | 34.58 | 65.42 |

**Figure 3:**
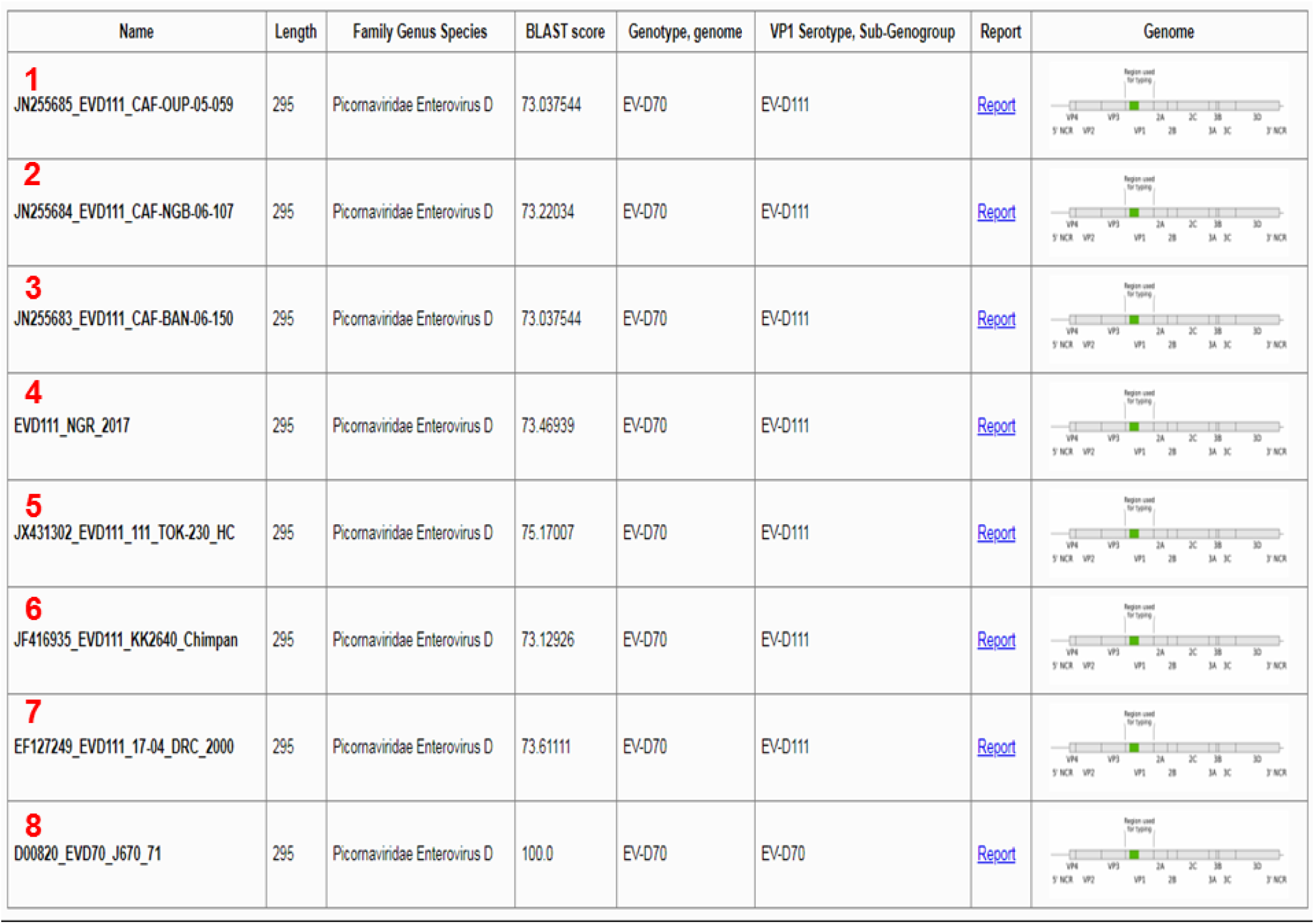
All the EV-D111 sequences in GenBank described till date alongside a reference EV- D70 and the EGT identification glitch.

### Phylogenetic Analysis

Phylogenetic analysis showed that two (sub-Saharan Africa [SSA] 1 and 2) lineages of E1 have been documented to circulate in sub-Saharan Africa from 1999 till date, and both have been detected in Nigeria (Figure 4). The E1 described in this study belongs to lineage SSA1 and seems to share a common ancestor with one of those we recovered (unpublished) on LLC-MK2 from sewage contaminated water in 2012 and another that was recovered from a child with AFP in Niger in 2014 (Fernandez-Garcia et al., 2017) (Figure 4).

**Figure 4:**
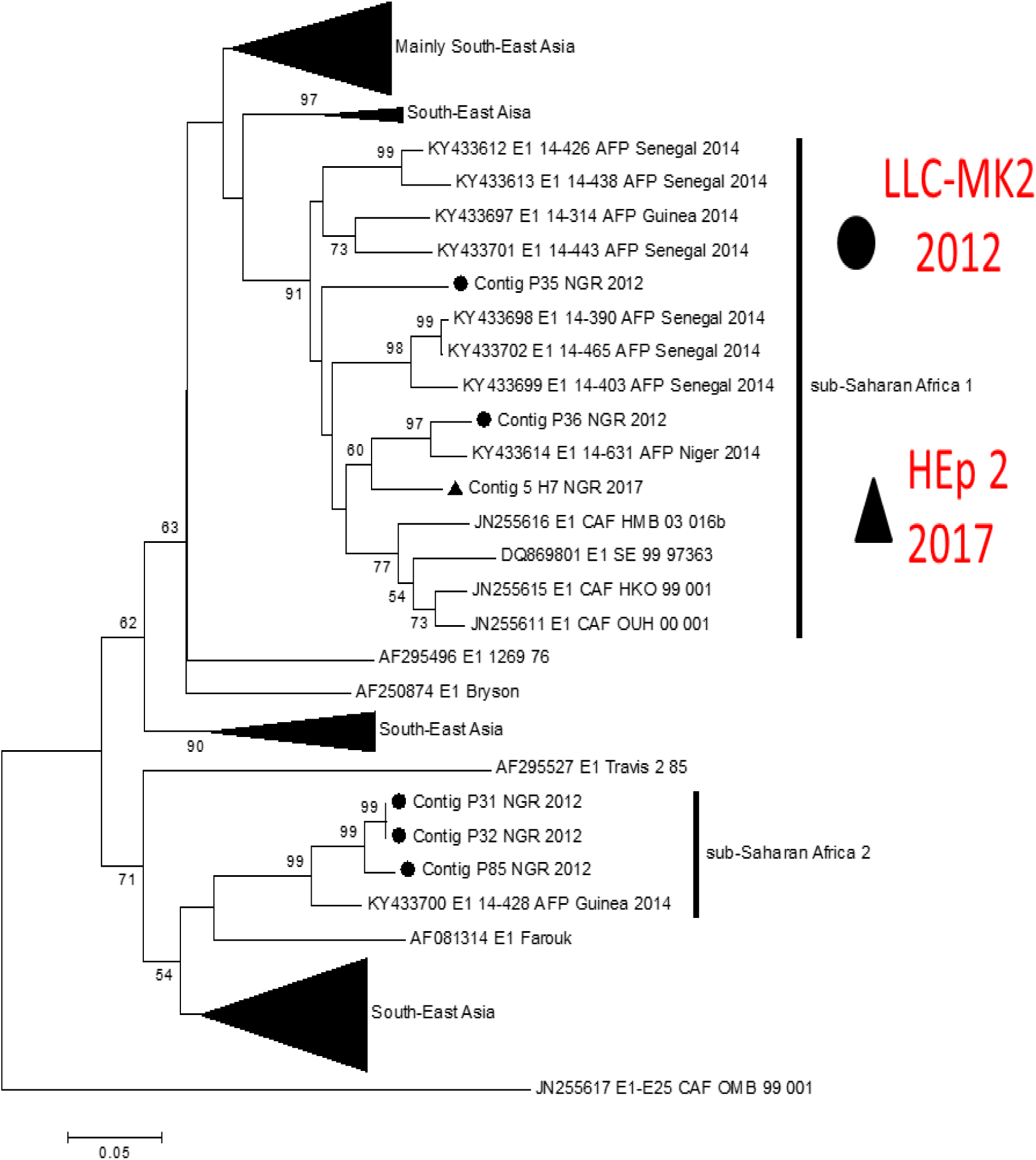
Phylogram of partial VP1 sequences of Echovirus 1. The phylogenetic tree is based on an alignment of partial E1 VP1 sequences available in GenBank and those generated by our group. The E1 described in this study is indicated with a black triangle. Other previously unpublished E1 sequences we generated in 2012 were added. Bootstrap values are indicated if >50%. Note that while E1 was isolated on Hep-2 in this study, all the E1s we isolated in 2012 were recovered on LLC-MK2 cell line.

Phylogenetic analysis also showed that two (1 & 2) lineages of EV-D111 have been documented globally (Figure 5). The EV-D111 lineage described in this study belongs to lineage 1 alongside those recovered in Cameroon and Central Africa Republic (CAF) (Figure 5). It is crucial to mention that, prior this study, only six (6) EV-D111 sequences have been described till date and all were detected in sub-Saharan Africa (Figure 6).

**Figure 5:**
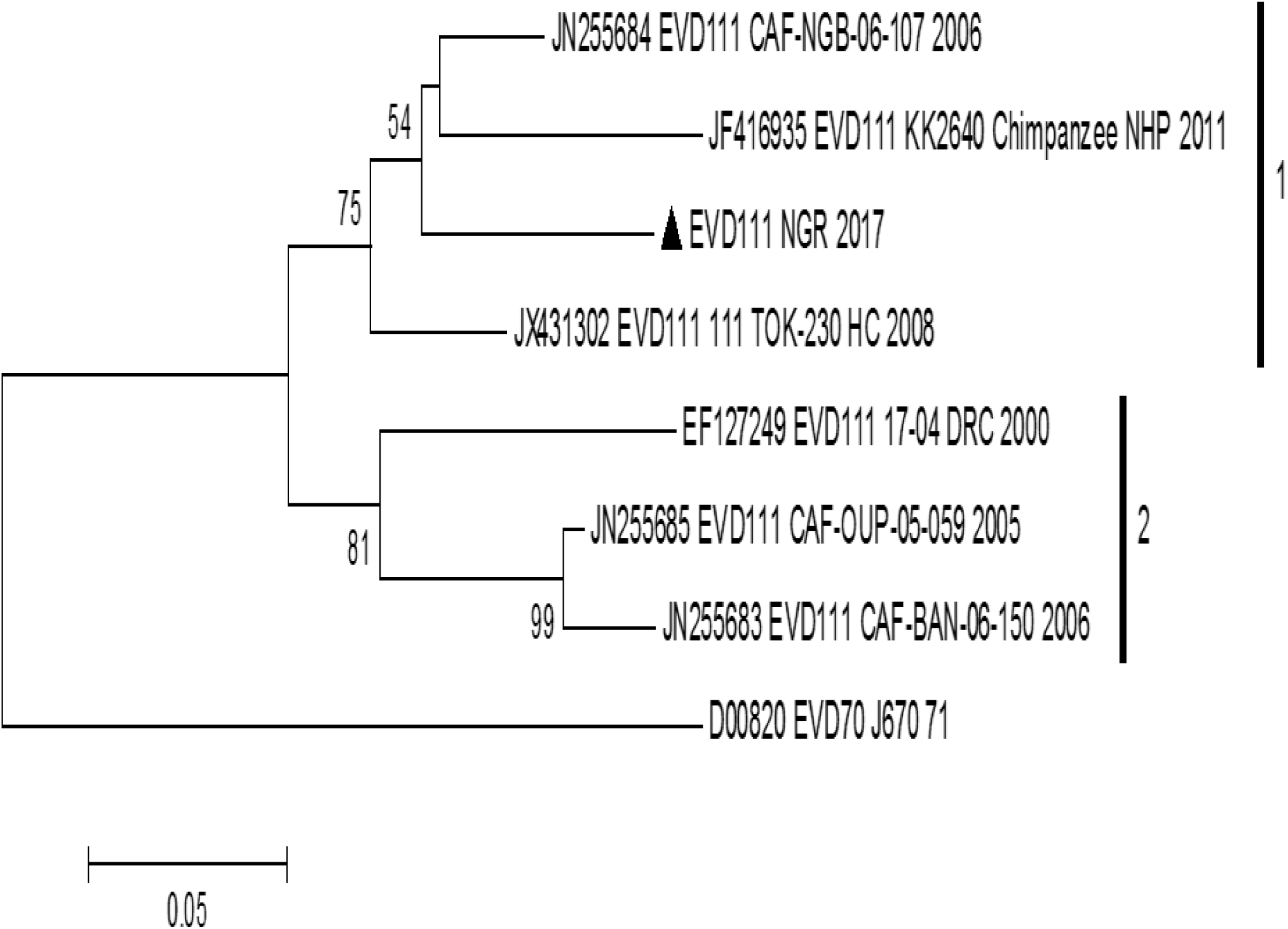
Phylogram of partial VP1 sequences of EV-D111. The phylogenetic tree is based on an alignment of partial EV-D111 VP1 sequences available in GenBank and the one described in this study. The EV-D111 described in this study is indicated with a black triangle. Bootstrap values are indicated if >50%.

**Figure 6:**
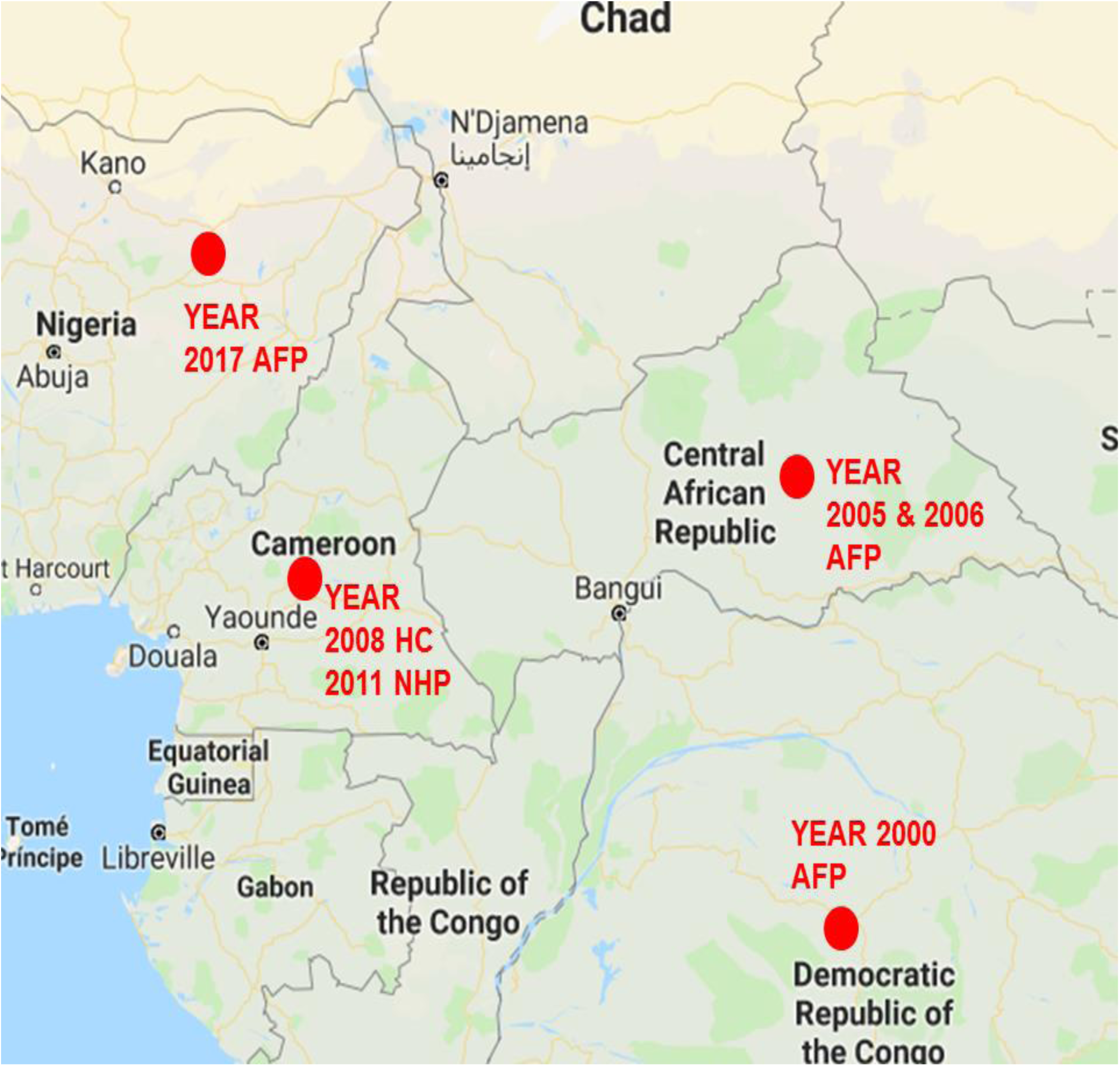
Global distribution of EV-D111 detection sources. It is crucial to note that, till date, all have been detected in sub-Saharan Africa.

## DISCUSSION

In this study we screened 59 CCSs with NR-CPE for Enteric Adenovirus or Group A Rotavirus and found them to be negative. Though, it is currently impossible to assay for nonspecific cytotoxicity and several other non-Enteric Adenoviruses and Reoviruses (asides the Group A Rotaviruses) remain unaccounted for in this study, the results suggest that cases of NR-CPE might not always be accounted for by Enteric Adenoviruses or Group A Rotaviruses.

We (Adeniji et al., 2018) had previously shown that EVs could be recovered from CCSs of NR- CPE by direct RT-snPCR (Nix et al., 2006, WHO, 2015). We (Adeniji et al., 2018) also showed that EVs were detectable by direct RT-snPCR in 30.8% (4/13; 2 each of EV-Cs and EV-Bs) of the CCSs. We have taken a step further to recover the EV-B and EV-C isolates on RD (Adewumi et al., 2017) and Hep-2 (Faleye et al., 2018b) cell lines, respectively. Considering, Hep-2 cell line supports replication of EV-Bs and has been documented to be better than RD in supporting the replication EV-Cs (Sadeuh-Mbab et al., 2013), we selected it for use in this study. Using Hep-2 cell line, we recovered four (4) isolates from the samples screened and showed unequivocally, that three of them were EVs. Of the EVs detected in this study, one (E1) belongs to EV-B while the remaining two are members of EV-D (EV-D111 and EV-D94). To our surprise, none of the isolates recovered were EV-C members despite the documented predilection of Hep-2 cell lines for EV-Cs (Sadeuh-Mba et al., 2013) and our previous findings of EV diversity in NR-CPE (Adeniji et al., 2018).

Specifically, 5% (3/59) of the NR-CPE CCSs screened in this study had EVs that were missed by the RD-L20b (R-L) cell-culture-based algorithm (WHO, 2003, 2004, 2015). We therefore confirm in this study that NR-CPE could sometimes be caused by EVs that possibly do not produce R-CPE in RD and L20b cell lines but do so in other cell lines like HEp-2. How often these EVs cause NR- CPE in R-L cell lines need further investigation. However, pending that, it seems 5% (this study) to 30% (Adeniji et al., 2018) of CCSs from NR-CPE on R-L cell lines might contain EVs.

Three EVs (E1, EV-D111 and EV-D94) were identified in this study. While VP1 data was available to identify E1 and EV-D111, the EV-D94 isolate had no VP1 data available (figure 2). It has been recommended (Harvala et al., 2018) and we have found it to be true (Adewumi et al., 2018) that in cases where VP1 data is not available, VP2 and VP4 can be utilized for EV identification. Hence, identification of the EV-D94 was done using a combination of the EGT and BLASTn result (data not shown). While the BLASTn result (data not shown) identified the isolate as EV-D94, the EGT could not assign it to any EV type using the VP2 and VP4 data. It however identified the isolate as EV-D94 using sequence data from the P3 genomic region, confirming the BLASTn result for the same region.

The EVs (E1, EV-D94 and EV-D111) described in this study (Table 2) are being described for the first time in Nigeria. In fact, should the information in GenBank and the Picornavirus Study Group website be real-time, as at the 4^th^ of November 2018, no complete genome of EV-D111 has been described and complete sequences exist for only the VP1 and VP4 genes. The EV-D111 sequence data we provide here, has the complete VP4, VP2, VP3, 2A, 2B, 2C, 3A and 3B genes (Figure 2). It also has partial sequences for 5-UTR, VP1 and 3C. Hence, some of the sequence data we provide for EV-D111 are being described for the first time globally. It is however, crucial to note that this valuable data would have been missed if the CCSs from these NR-CPEs had not been re-screened. We believe, that as the EV community move more towards direct detection of EVs from samples using RT-PCR and NextGen sequencing strategies (Isaacs et al., 2018, Joffret et al., 2018, Majumdar and Martin, 2018), it is likely that we have less of these omissions. However, as we have previously shown (Adeniji et al., 2017), it should be noted that RT-PCR assays have limit of detection and their sensitivity can be enhanced when augmented with cell culture.

We (Adeniji and Faleye, 2014, Faleye and Adeniji, 2015, Adeniji et al., 2017) have previously shown how choice of cell line and the consequent cell culture bias influences our perception of the diversity landscape of EVs in any region. The findings of this study further illuminate this observation. The E1 described in this study was isolated on Hep-2 cell line (Figure 3). During our (Adeniji and Faleye, 2015) studies on sewage samples in 2012, we had previously detected E1 (unpublished) on LLC-MK2 cell line and phylogenetic analysis showed that one of the E1s recovered from sewage and isolated on LLC-MK2 in 2012 shares a common ancestor with the strain described in this study (figure 3). Thus, if these cell lines were not included, it would have been assumed that E1 was neither present nor circulating in Nigeria. This further exemplifies the impact of cell culture bias on our perception of EV diversity in any region. Hence, any study in which cell culture is the basis for defining EV diversity in any population or sample set should be interpreted with the understanding that what the study documents is only what the bias permits. It is therefore crucial to mention that expanding the combination of cell lines used for enterovirus detection can and really do help expand the range of EV types recoverable in cell culture.

To the best of our knowledge, till date, EV-D111 has only been detected in sub-Saharan Africa (Juntila et al., 2007, Harvala et al., 2011, Bessaud et al., 2012, Sadeuh-Mba et al., 2013). Why this is the case is not clear. It therefore remains to be shown whether EV-D111 is present or circulating outside sub-Saharan Africa. Pending that, it seems EV-D111 is confined to the region (Figure 5). Though confined to sub-Saharan Africa, EV-D111 has been recovered from feaces of NHPs (Harvala et al., 2011), healthy children (Sadeuh-Mba et al., 2013) and those with AFP (Juntila et al., 2007, Bessaud et al., 2012, this study). Hence, its zoonotic potential is clear. More importantly, the strain found in the AFP case in Nigeria (this study, Figure 4) belongs to lineage 1; the same lineage to which that found in NHP in Cameroon (a country sharing border with Nigeria) belongs. Considering the global distribution of EV-D68 (another member of EV species D) and its recurrent association with outbreaks with severe clinical manifestations (Esposito et al., 2015, Knoester et al., 2018), it is essential that the molecular epidemiology of this EV type be better investigated.

Currently, it seems the EGT is finding it difficult to type EV-D111. We first observed this with the EV-D111 isolate described in this study (Figure 2). We consequently, checked all the six (6) EV-D111 strains described and available in GenBank prior this study and confirmed our observation (Figure 6). To be precise, using the capsid region data (P1), the EGT identifies EV- D111 strains as EV-D70 genotype and EV-D111 serological type (Figure 6). It however does no such thing with E1 (Figure 2). As at the time of writing this manuscript, EV-D70 and EV-D111 remain as independent EV types on the EV-D page of the Picornavirus Study Group website (http://www.picornaviridae.com/enterovirus/ev-d/ev-d.htm). It is therefore essential that the EGT software is checked to fix this glitch.

## ACKNOWLEDGEMENTS

We thank the WHO National Polio Laboratory in Ibadan, Nigeria for providing the HEp-2 cell line and anonymous cell culture supernatants analyzed in this study. We thank Oladipo Kolawole Elijah for providing us reference Adenovirus 1 and 6 from BBSRC. This study was partly funded by a TETFund grant to Adeniji J.A. PhD.

